# ProteomeGenerator: A framework for comprehensive proteomics based on de novo transcriptome assembly and high-accuracy peptide mass spectral matching

**DOI:** 10.1101/236844

**Authors:** Paolo Cifani, Avantika Dhabaria, Akihide Yoshimi, Omar Abdel-Wahab, John T. Poirier, Alex Kentsis

**Affiliations:** Molecular Pharmacology Program, Sloan Kettering Institute, Memorial Sloan Kettering Cancer Center, New York, NY.; Human Oncology and Pathogenesis Program, Memorial Sloan Kettering Cancer Center New York, NY.; Department of Medicine, Memorial Sloan Kettering Cancer Center, New York, NY.; Department of Pediatrics, Memorial Sloan Kettering Cancer Center, New York, NY.; Departments of Pediatrics, Pharmacology, and Physiology & Biophysics, Weill Cornell Medical College, Cornell University, New York, NY.

## Abstract

Modern mass spectrometry now permits genome-scale and quantitative measurements of biological proteomes. However, analyses of specific specimens are currently hindered by the incomplete representation of biological variability of protein sequences in canonical reference proteomes, and the technical demands for their construction. Here, we report ProteomeGenerator, a framework for *de novo* and reference-assisted proteogenomic database construction and analysis based on sample-specific transcriptome sequencing and high-resolution and high-accuracy mass spectrometry proteomics. This enables assembly of proteomes encoded by actively transcribed genes, including sample-specific protein isoforms resulting from non-canonical mRNA transcription, splicing, or editing. To improve the accuracy of protein isoform identification in non-canonical proteomes, ProteomeGenerator relies on statistical target-decoy database matching augmented with spectral-match calibrated sample-specific controls. We applied this method for the proteogenomic discovery of splicing factor *SRSF2*-mutant leukemia cells, demonstrating high-confidence identification of non-canonical protein isoforms arising from alternative transcriptional start sites, intron retention, and cryptic exon splicing, as well as improved accuracy of genome-scale proteome discovery. Additionally, we report proteogenomic performance metrics for the current state-of-the-art implementations of SEQUEST HT, Proteome Discoverer, MaxQuant, Byonic, and PEAKS mass spectral analysis algorithms. Finally, ProteomeGenerator is implemented as a Snakemake workflow, enabling open, scalable, and facile discovery of sample-specific, non-canonical and neomorphic biological proteomes (https://github.com/jtpoirier/proteomegenerator).

## INTRODUCTION

Functional analysis of physiologic and pathologic cell activities requires accurate and complete identification and quantification of all involved effector molecules. Such global studies are principally based on the decoding and assembly of the human genome [1,2]. Recent advances in messenger RNA (mRNA) sequencing and bioinformatics now enable routine analysis of biological gene expression [3-5]. However, direct and proteome-wide studies of proteins and their biological variation remain confined to specialized approaches [6-8].

Modern mass spectrometry now permits genome-scale and quantitative measurements of biological proteomes [9-13]. This approach is based on mass spectrometric analysis of peptides, generated by proteolysis of biological proteomes, followed by matching their observed fragmentation spectra to those predicted from known or expected protein sequences [14,15]. Generally, this is accomplished using statistical peptide-spectrum matching techniques that leverage scoring functions to assess the similarity of observed and theoretical mass spectra [16], with the corresponding confidence of spectral identification expressed as a global false discovery rate (FDR), estimated using target-decoy approaches [17].

Peptide identification using peptide-spectrum matching (PSM) and target-decoy FDR estimation is based on the fundamental assumption that mass spectrometry search databases contain a complete and accurate list of all potential protein sequences. Consequently, sensitivity and specificity of peptide identification depend on the fidelity of the target and decoy databases. Advances in genome analysis and assembly have led to the development of high-quality databases of consensus protein sequences such as UniProt and RefSeq [1,2,18,19]. However, germline and somatic genetic variants, messenger RNA (mRNA) splicing, and other biological processes can diversify polypeptide sequences, thereby generating sequence variants that are not catalogued in the consensus or canonical databases [20,21]. These sources of proteome variation are particularly prevalent in human cancers, which can be caused by structural aberrations in genes and dysregulation of their expression [22-24], ultimately hindering the discovery of cancer proteomes based on reference consensus databases.

In principle, proteogenomic approaches that integrate genome and transcriptome sequencing data with mass spectrometric protein analysis can overcome this limitation, by generating sample-specific target databases for proteomic analysis that more accurately reflect the expressed proteome [25-31]. Such an approach was first introduced to support gene annotation using proteomic data [32], and has since become a powerful tool for quantitative and integrative studies [30,33-38]. In addition, related approaches were recently developed for cancer biology [39-44], and immunology studies [45]. Specifically, sequences of expressed gene transcripts obtained by high-throughput sequencing of mRNA (RNA-seq) are a convenient source for proteogenomic sample-specific database construction for two main reasons: i) these measurements reflect sequence variability introduced by transcriptional and post-transcriptional processes, and ii) restriction of the mass spectral match search space to the specifically expressed proteins can improve its sensitivity and accuracy, particularly as compared to proteins predicted from translation of all possible reading frames [46].

Based on this rationale, several approaches have recently been developed to generate customized mass spectrometry search databases from RNA-seq data [28,44,47,49]. However, sensitivity and accuracy of proteogenomic detection of neomorphic and non-canonical proteins remain limited by the under-sampling of rare peptides and challenges in the generation of sample-specific databases from non-strand-specific short-read RNA-seq data [25,27]. Furthermore, while reference sequence databases such as UniProt are manually curated to improve accuracy [18], automated workflows are needed to enable facile sample-specific proteogenomic analyses at scale.

Here, we describe ProteomeGenerator, an open, modular, and scalable framework for *de novo* and referenced proteogenomic database construction and analysis written in the Snakemake workflow management system. To improve the accuracy of peptide-spectrum matching, we augmented established statistical target-decoy mass spectral matching with spectral-match calibrated sample-specific controls and parameterized this method using four current state-of-the-art mass spectrometry search algorithms. Lastly, we used ProteomeGenerator for genome-scale proteomic discovery of splicing factor-mutant leukemia cells, based on the integration of deep mRNA sequencing, and multidimensional, high-capacity nano-scale chromatography and high-accuracy mass spectrometry. This led to the high-confidence identification of non-canonical protein isoforms arising from alternative transcription start sites, intron retention, and cryptic exon splicing, as well as improved accuracy of genome-scale proteome discovery as compared to conventional approaches.

## EXPERIMENTAL PROCEDURES

### Reagents

Mass spectrometry grade (Optima LC/MS) water, acetonitrile (ACN), and methanol were from Fisher Scientific (Fair Lawn, NJ). Formic acid of >99% purity (FA) was obtained from Thermo Scientific. All other reagents at MS-grade purity were obtained from Sigma-Aldrich (St. Louis, MO).

### Cell culture

Human K052 cells were obtained from the Japanese Collection of Research Bioresources Cell Bank, identity confirmed using STR genotyping (Genetica DNA Laboratories, Burlington, NC), and cultured as described [50]. Cells were collected while in exponential growth phase, washed twice in ice-cold PBS, snap frozen and stored at −80°C. Protein extraction and proteolysis was performed as previously described [51]. Briefly, frozen cell pellets were thawed on ice, resuspended in 6 M guanidinium hydrochloride, 100 mM ammonium bicarbonate at pH 7.6 (ABC), and lysed using the E210 adaptive focused sonicator (Covaris, Woburn, CA). The protein content in cell lysate was determined using the BCA assay, according to the manufacturer’s instructions (Pierce, Rockford, IL). Upon reduction and alkylation, proteomes were digested using 1:100 w/w (protease:proteome) LysC endopeptidase (Wako Chemical, Richmond, VA) and 1:50 w/w MS sequencing-grade modified trypsin (Promega, Madison WI). Digestion was stopped by acidifying the reactions to pH 3 using formic acid (Thermo Scientific), and peptides were subsequently desalted using solid phase extraction using C18 Macro Spin columns (Nest Group, Southborough, MA).

### mRNA sequencing

RNA was extracted using QIAGEN RNeasy columns (Qiagen, Valencia CA). Poly(A)-selected, unstranded Illumina libraries were prepared with a modified TruSeq protocol, and 0.5x AMPure XP beads (Beckman Coulter, Indianapolis IA) were added to the sample library to select for fragments of <400 bp, followed by 1x beads to select for fragments of >100 bp. These fragments were then amplified with PCR (15 cycles) and sequenced on the Illumina HiSeq 2000 (100 million 2 × 49 bp reads/sample).

### Proteome Generator

ProteomeGenerator is written in Snakemake, a scalable, Python-based workflow management system (Figure 2) [52]. The entire workflow is available for download at https://github.com/jtpoirier/proteomegenerator. ProteomeGenerator ingests RNAseq data, which is aligned to a reference genome by the STAR splice aware aligner (v. 2.5.2a) [53]. Aligned reads are subsequently filtered to exclude low quality and poorly mapping reads using samtools (v. 1.3) [54]. A sample-specific transcript model is then assembled either *de novo* or with assistance from reference transcript model if one is available using StringTie (v. 1.3.3b) [55], with simultaneous filtering for transcripts with coverage ≥2.5, length ≥300 base pairs, and abundance of ≥1% of expressed transcripts for a given gene. All resulting transcript models are then merged with StringTie --merge using an expression threshold of 1 transcript per million with permissive intron inclusion. The resulting merged transcript model is then used to generate corresponding cDNA sequences using gffread (v. 0.9.8). The longest uninterrupted open reading frame is detected within each cDNA using TransDecoder (v. 2.1, https://github.com/TransDecoder) [56]. Shorter open reading frames with low expect values when searched against the UniProt database using BLAST (v. 2.2.31) are retained [57]. The predicted longest open reading frames are subsequently mapped back to genomic coordinates and translated to their respective unique peptide sequences.

### Peptide and database characterization

Peptides assigned to mass spectra are mapped back to their genomic coordinates using ProteomeGenerator for visualization in the Integrative Genomics Viewer [58]. Unique tryptic peptide search space was calculated for each database using the EMBOSS [59] tool pepdigest and filtered to include all peptides of at least 6 amino acids in length having a molecular weight between 600 and 4000 Daltons.

### Databases

Consensus protein sequences databases were downloaded from UniProt [18] as of January 2016 (*Homo sapiens*), September 2015 (*Archaebacteria loki*) and June 2014 (*Escherichia coli*). Contaminant sequences were retrieved from cRAP [60] as of June 2014.

### High-resolution peptide chromatography

Peptide chromatographic fractionation was performed using the Alliance e2695 high-performance liquid chromatograph (Waters, Milford MA). High-pH reverse phase separation was performed using the Xselect CSH 3.0 mm × 150 mm column (Waters, part no. 186006728) at constant flow-rate of 250 μl/min. After an initial equilibration at 100% buffer A (50 mM ammonium hydroxide in water, pH 10) for 5 minutes, peptides were resolved by a 75 minutes 0–70% gradient of buffer B (80% ACN in water, pH 10), followed by 10 minutes at 100% buffer B. The eluate was collected in 0.5 ml aliquots using a fraction collector between 25 and 75 minutes, lyophilized to dryness in a vacuum centrifuge, and stored at −80°C until analysis. Before LC-MS analysis, peptides were resuspended in 20 μl 0.1% formic acid in water, and 2 μl were analyzed.

Strong cation exchange chromatography was performed using the Xselect Hi Res SP, 7 μm, 4.6 mm × 100 mm column (Waters, part no. 186004930) at constant flow-rate of 500 μl/min. After an initial equilibration at 100% buffer A (0.1 % formic acid, 5% ACN) for 5 minutes, peptides were resolved by a 80 minutes 0–30% gradient of buffer B (1 M KCl, 5% ACN), followed by 5 minutes gradient 30–50% buffer B, and a final hold for 5 minutes at 100% buffer B. The eluate was collected in 1 ml aliquots using a fraction collector between 25 and 85 minutes, and lyophilized to dryness in a vacuum centrifuge. Pellets were desalted by solid phase extraction using C18 Macro Spin columns, and stored at −80°C until analysis. Before LC-MS analysis, peptides were resuspended in 20 μl 0.1% formic acid in water, and 2 μl were analyzed.

### Nanoscale liquid chromatography

Nanoscale liquid chromatography experiments were performed using the Ekspert NanoLC 425 chromatograph (Eksigent, Redwood city, CA), equipped with an autosampler module, two 10-port and one 6-port rotary valves, and one isocratic and two binary pumps. Column and emitter fabrication were performed as previously described [61]. Briefly, samples were initially aspirated into a 10 μl PEEK sample loop. Chromatographic columns were fabricated by pressure filling the stationary phase into silica capillaries fritted with K-silicate. Reversed phase columns were fabricated by packing Reprosil 1.9 μm silica C18 particles (Dr. Meisch, Ammerbauch-Entrigen, Germany) into 75 μm x 40 cm fritted capillaries. Trap columns were fabricated by packing Poros R2 10 μm C18 particles (Life Technologies, Norwalk, CT) into 150 μm x 4 cm fritted capillaries. Vented trap-elute architecture was used for chromatography [62]. Peptides were resolved by reversed phase chromatography hyphenated to the nano-electrospray ion source. Upon valve switch to connect the trap column in line with the analytical reversed phase column and ion emitter, the pressure was equilibrated at a flow of 250 nl/min for 5 minutes in 5% buffer B (ACN, 0.1% FA) in buffer A (water, 0.1% FA). Subsequently, a 120-minutes (high-pH reverse phase samples) or 180 minutes (SCX samples) linear gradient of 5-40% of buffer B was used to resolve peptides, followed by a 5 minutes 40-80% gradient prior to column wash at 80% buffer B for 30 minutes.

### Nano-electrospray ionization and Orbitrap mass spectrometry

Electrospray emitters with terminal opening diameter of 10 μm were obtained from New Objective (Woburn, MA). The emitter was connected to the outlet of the reversed phase column using a metal union that also served as the electrospray current electrode. Electrospray ionization was achieved using constant 1700 V voltage. During column loading, the electrospray emitter was washed with 50% aqueous methanol using the DPV-565 PicoView ion source (New Objective).

For all measurements, we used the Orbitrap Fusion mass spectrometer (Thermo Scientific, San Jose, CA). Precursor scans in the 400-2000 Th were performed in the orbitrap detector at 120,000 resolution, with 100 ms maximum injection time, and automatic gain control set at 10^5^ ions. Fragment spectra were recorded in the linear ion trap in rapid mode, with maximum injection time of 75 ms and target of 10^4^ ions, and 1.2 Da quadrupolar precursor selection.

### Data Analysis

Peptide-spectral matching calculations were performed using a custom-built computer server equipped with 4 Intel Xeon E5-4620 8-core CPUs operating at 2.2 GHz, and 512 GB physical memory (Exxact Corporation, Freemont, CA). Peptide-spectral matching was performed using SEQUEST HT and Percolator [16,63,64] as part of Proteome Discoverer 2.1.0.81 (Thermo Scientific), Byonic 2.7.84 [65,66], MaxQuant 1.5.4.1 [67,68], and PEAKS 8.0 [69]. For all searches, MS1 and HCD MS2 mass tolerances were set to 10 ppm and 0.6 Da, respectively. Cysteine carbamidomethylation was set as fixed, while methionine oxidation and glutamine and asparagine deamidation were set as variable modifications, with a maximum of 3 variable modifications per peptide. Only peptides containing 7–35 residues and up to 2 missed trypsin cleavages were considered. Tryptic peptides observable by mass spectrometry were predicted based on proteome composition using the generate-peptides utility of Crux [70].

### Data availability

ProteomeGenerator is openly available at https://github.com/jtpoirier/proteomegenerator. RNA sequencing data are available from the Gene Expression Omnibus. Mass spectrometry raw files and search results are publicly available through the ProteomeXchange data repository with identifiers PXD008381 and PXD008463 [71].

## RESULTS

### ProteomeGenerator for deep genome-scale transcriptomic and proteomic integration

Consensus reference proteomes provide a useful set of canonical protein sequences to be used for statistical peptide-spectral matching in mass spectrometry proteomics. However, these databases do not fully recapitulate the natural and pathologic variation in protein sequences, therefore limiting the sensitivity and specificity of proteome discovery. To enable facile construction of sample-specific proteomic databases containing expressed non-canonical protein variants, we developed a proteogenomic framework, termed ProteomeGenerator, programmed in the Snakemake workflow management system for open and scalable analysis. This method i) generates a sample-specific protein sequence database based on de novo or referenced transcriptome assembly using high-coverage RNA-seq data, ii) deep mass spectrometric analysis of the matched proteome using high-resolution nanoscale chromatography and high-accuracy mass spectrometry, and iii) accurate protein isoform identification using statistical target-decoy database matching augmented with spectrally calibrated sample-specific controls (Figure 1).

**Figure 1.**
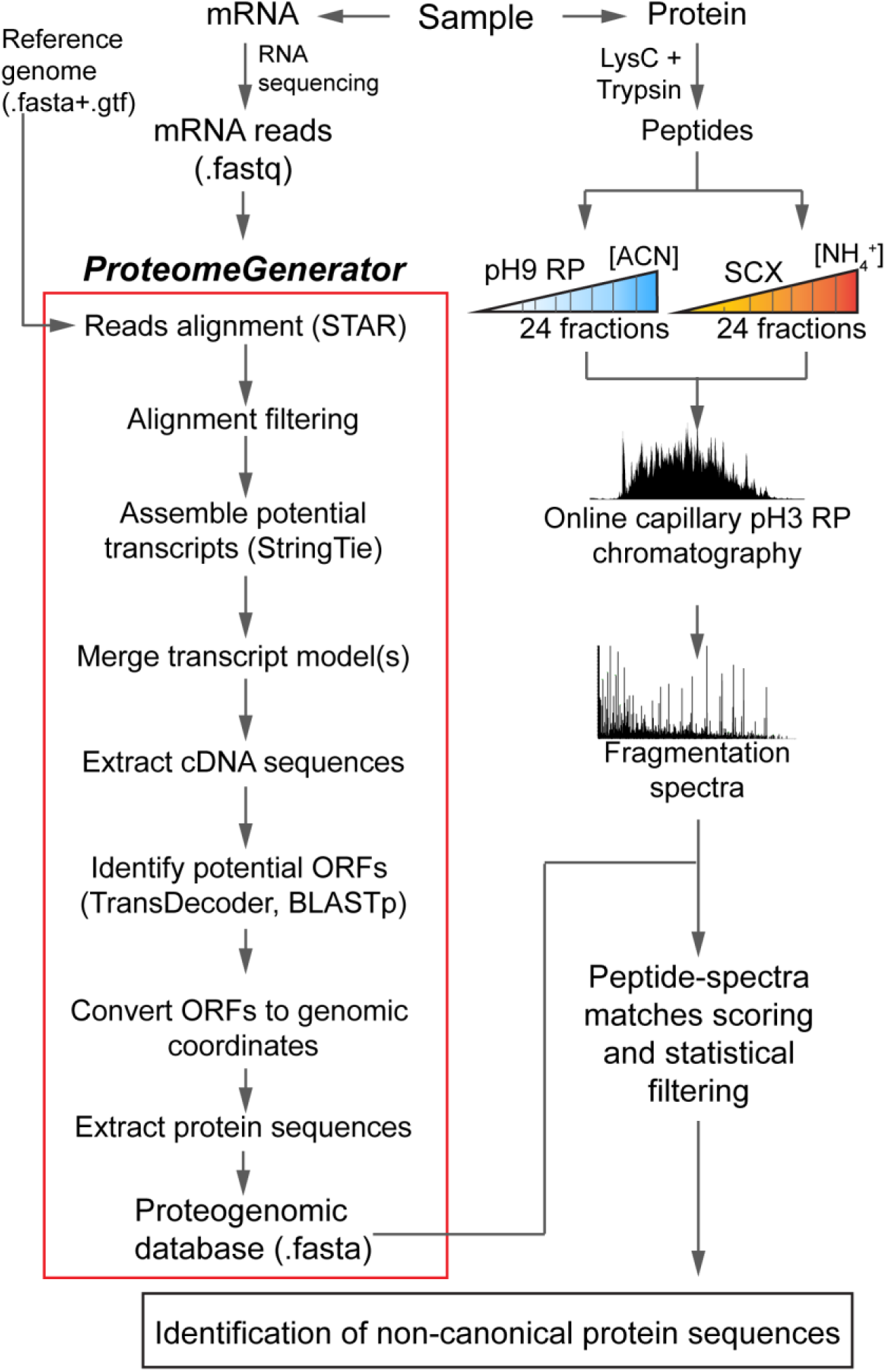
ProteomeGenerator overview. Transcriptomes and proteomes from the same biologic sample are analyzed in parallel by high-coverage Illumina sequencing and high-resolution, high-accuracy mass spectrometry, respectively. ProteomeGenerator assembles fastq-formatted mRNA sequencing reads into predicted transcripts, identifies reading frames and isoforms, and produces fasta-formatted proteogenomic (PGX) databases containing canonical and non-canonical expressed protein isoforms for subsequent mass spectrometry searches.

We applied this method to analyze the human K052 leukemia cells harboring splicing factor *SRSF2* mutations, which were recently described to cause recurrent mRNA mis-splicing, and are therefore expected to express non-canonical and neomorphic protein isoforms [50]. Thus, we extracted mRNA and obtained high-coverage RNA-seq data of nearly 60 million reads by Illumina sequencing. We then used ProteomeGenerator to process the raw sequencing reads in the following steps: i) two-pass splice aware alignment to the user-supplied reference genome, in this case GRCh38; ii) assembly of possible transcript isoforms using StringTie either de novo or assisted by the user-supplied transcript model, in this case GENCODE v20; iii) prediction of the longest open reading frame from for each possible transcript; and iv) generation of fasta-formatted proteogenomic database comprising unique protein sequences for subsequent peptide sequence-mass spectral matching (Figure 2).

**Figure 2.**
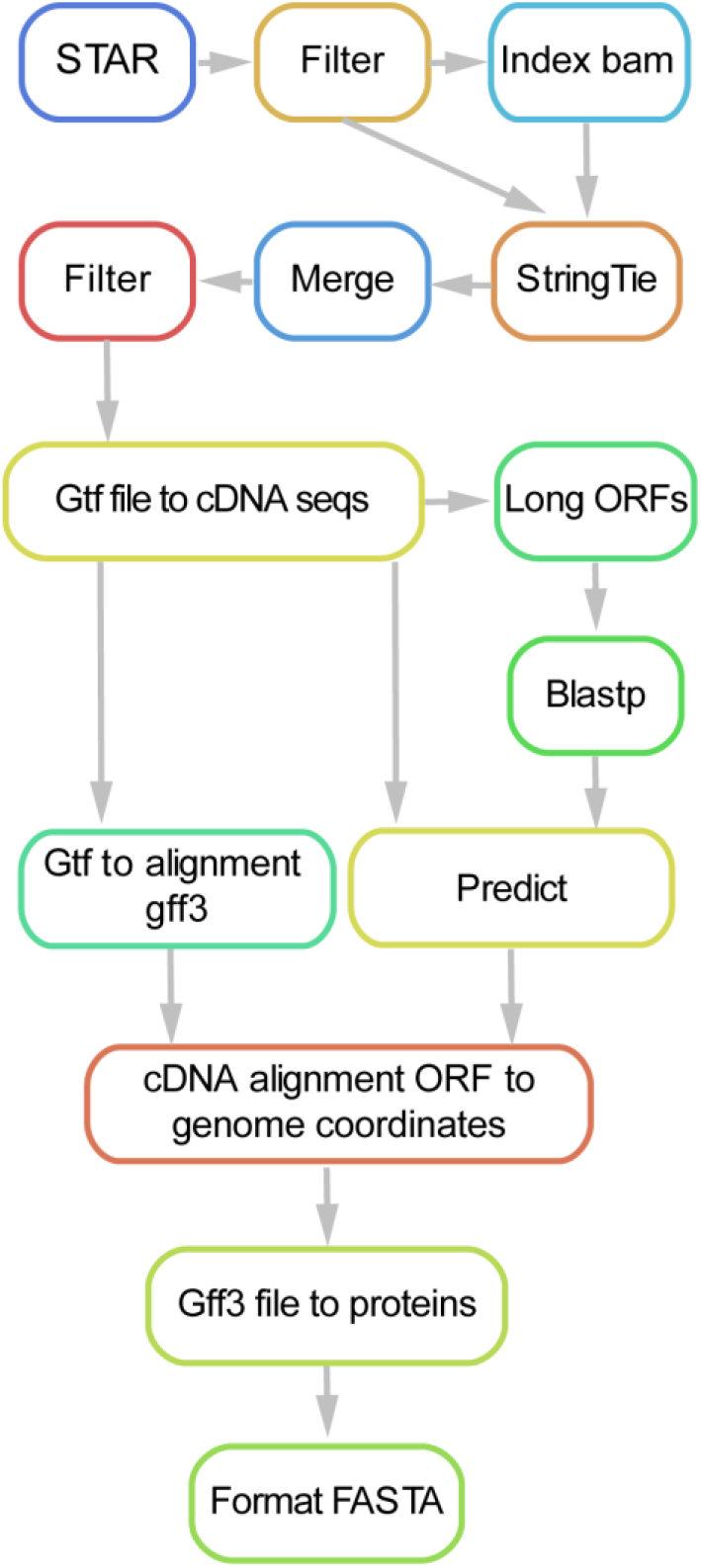
Schema for the ProteomeGenerator snakemake workflow. Sequencing reads are aligned using STAR, followed by their de novo and referenced assembly intro transcriptomes using StringTie, and processing to identify reading frames and protein isoforms.

In the case of analyzed K052 cells, the constructed proteogenomic database, to which we refer as PGX, consisted of 17,348 protein entries, expected to produce 743,148 observable tryptic peptides with lengths between 7 and 35 residues, assuming 1 missed trypsin cleavage (Figure 3). For comparison, the canonical reference human proteome in the UniProt database (as of March 2016), contained 42,123 proteins corresponding to 1,460,257 mass spectrometry observable tryptic peptides. The PGX database contained 37,158 peptides with no counterparts in UniProt, presumably originating from novel predicted protein isoforms specific for splicing factor-mutant K052 cells and consequently not annotated in UniProt. In addition, the PGX database was approximately 51% smaller with respect to mass spectrometry observable peptides as compared to UniProt, presumably because it contained only expressed protein isoforms, in contrast to the canonical reference proteomes, which contain the protein complement of all canonical protein isoforms.

**Figure 3.**
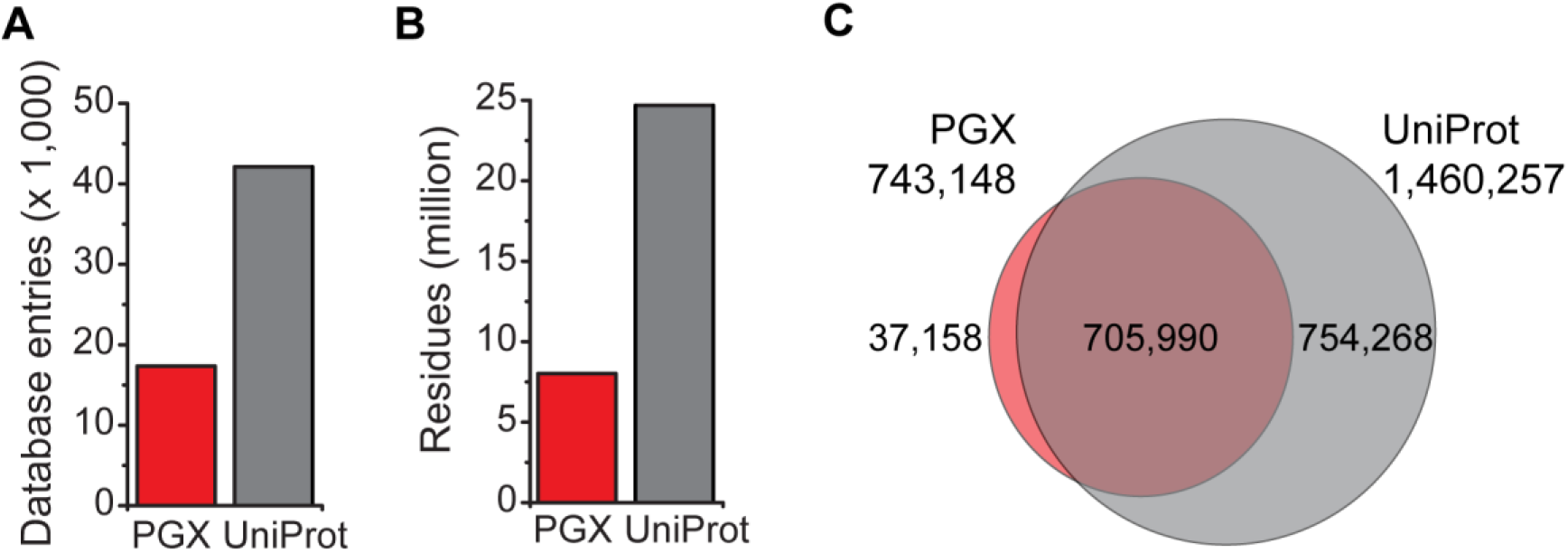
Comparison of the canonical and proteogenomic protein databases, displaying (A) number of protein entries, (B) number of amino acid residues, and (C) theoretical tryptic peptides amenable for mass spectrometry analysis specific for either UniProt, PGX, or common to both.

### Genome-scale mass spectrometry proteomics

To generate mass spectrometry data of sufficient coverage to sample genome-scale proteomes, we used recently developed high-resolution, multi-dimensional nanoscale chromatography and high-accuracy mass Orbitrap spectrometry. To increase chromatographic resolution, we leveraged the orthogonal properties of strong-cation exchange and alkaline reversed phase as compared to acidic reversed phase chromatography, thereby enabling the detection of low abundance and rare peptides and their isoforms [72]. Each peptide fraction was then resolved by high-resolution nano-scale reverse phase chromatography and analyzed by nanoelectrospray ionization mass spectrometry, and nearly 3 million high-resolution precursor and fragmentation spectra were recorded.

To assess sampling efficiency, we used statistical database matching against UniProt to identify unique peptides and proteins at 1% FDR. Using a sub-sampling analysis, we observed that this strategy indeed maximized the sensitivity of detection of canonical proteins, though peptide sampling was apparently incompletely saturated (Supplementary Figure 1A).

**Supplementary Figure 1.**
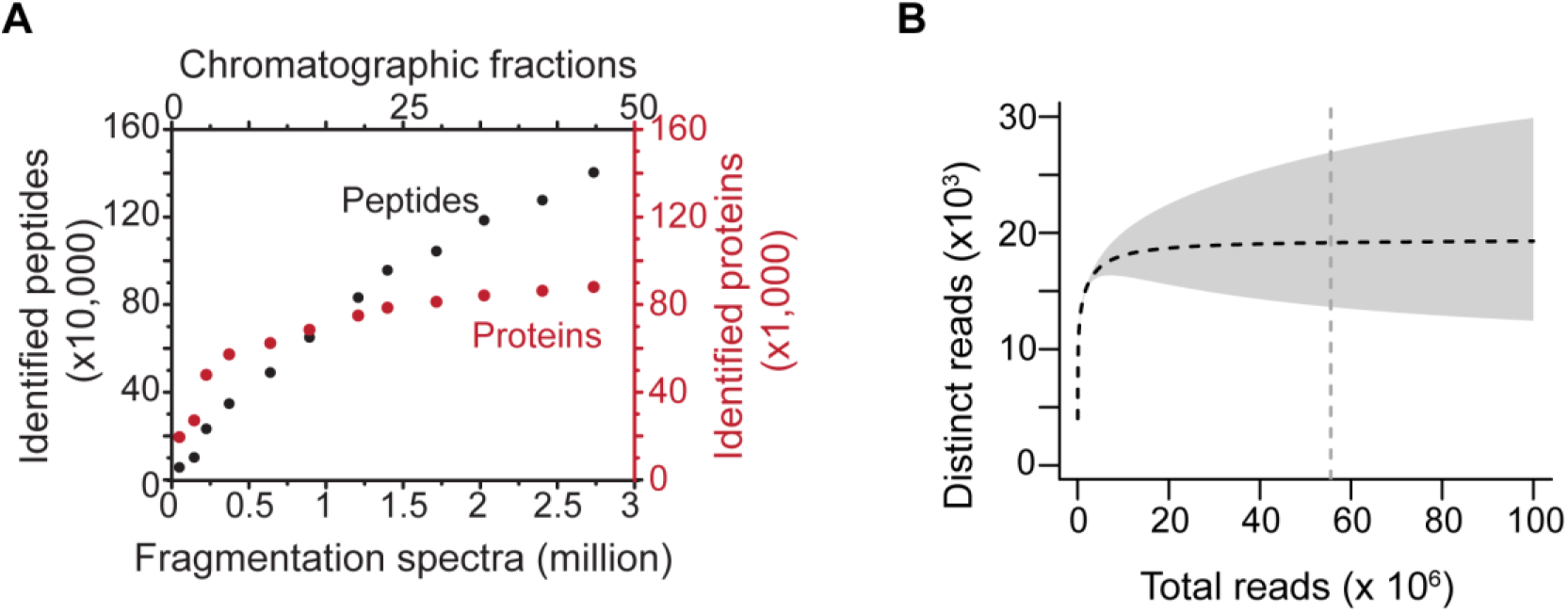
Subsampling analysis of mass spectrometric (A) RNA sequencing (B) dataset. Plateauing of the number of sequences detected indicates reaching of the limit of detection for the specific analytical method used, as observed in transcriptomic (B) but not in proteomic (A) datasets.

Likewise, we found that high-coverage mRNA sequencing apparently saturated sampling of unique sequence reads obtained by transcriptomic analysis (Supplementary Figure 1B). Thus, high-resolution chromatography c oupled with high-accuracy mass spectrometry and high-coverage transcriptome sequencing is suitable for genome-scale proteogenomics.

### Accurate proteome discovery using statistical target-decoy matching with spectral-match calibration

Having obtained genome-scale mass spectrometry data and the transcriptome-assembled PGX database based on high-coverage mRNA sequencing, we next sought to identify algorithms for scoring peptide-spectral matches and estimating FDR confidence suitable for proteogenomic analysis. Such algorithms should ideally not only maximize sensitivity (i.e. the fraction of identified spectra), but also ensure high specificity, in particular when searching a non-curated proteogenomic target database potentially containing erroneous sequences. To calibrate the sensitivity and specificity of these algorithms, we introduced two negative controls [73,74]: i) a set of mass spectra from a non-human proteome (*E. coli*) recorded under identical experimental conditions as the experimental human proteome, and ii) a set of protein sequences from evolutionarily divergent non-human species with minimal identity to the experimental human proteome, in this case the recently published proteome of *A. loki* archaebacteria [75], as confirmed by direct comparisons (Figure 4A–B).

**Figure 4.**
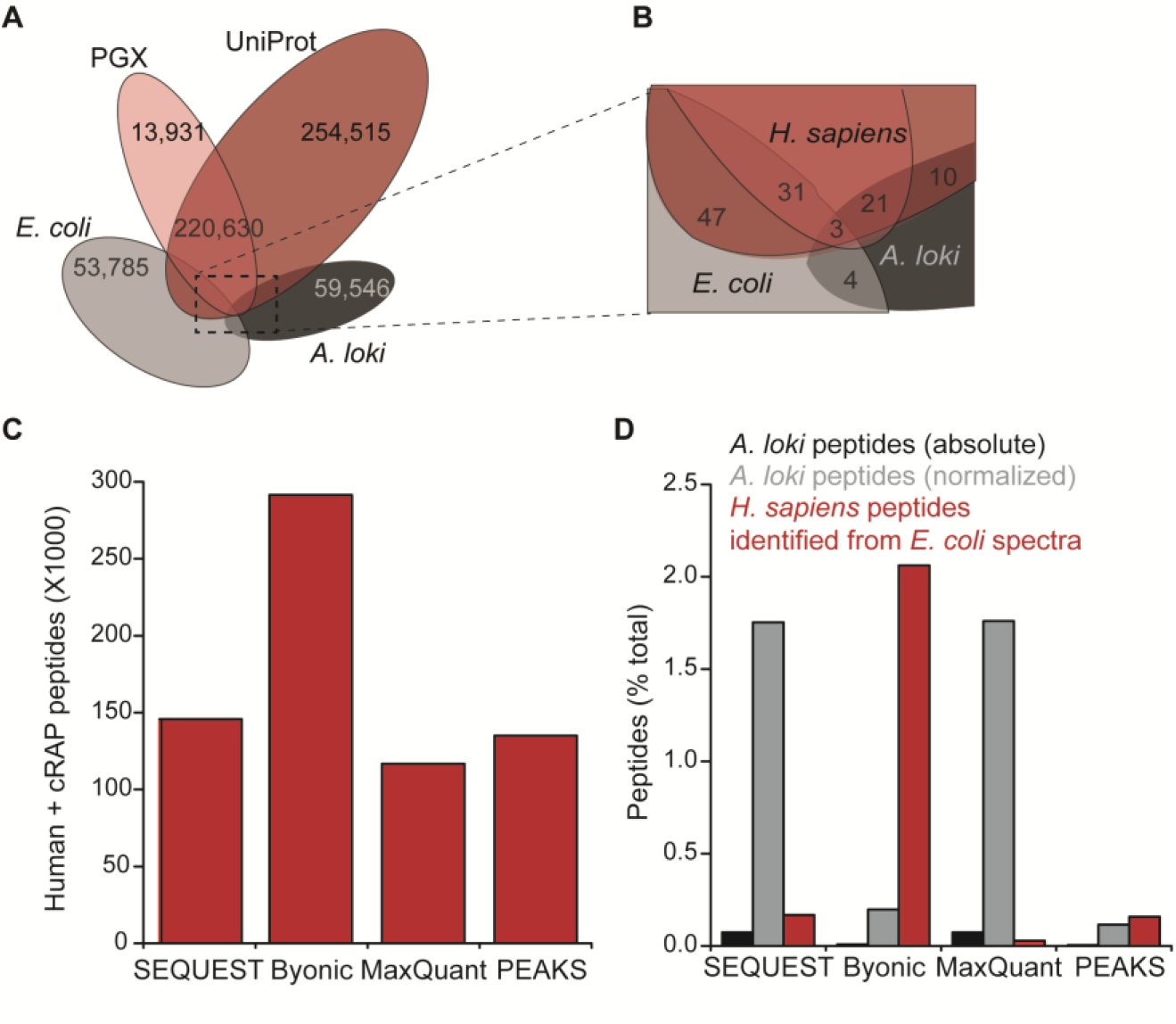
Sensitivity and specificity of mass spectrometry search algorithms. (A, B) Comparison of unique theoretical peptides in the experimental PGX proteome, canonical UniProt, and bacterial proteomes used as negative controls. (C) Sensitivity of tested algorithms, expressed as the number of identified peptides. (D) Specificity of tested algorithms, evaluated from the fraction of peptide-spectrum matches mapped to the negative controls. The PSM fraction mapped to *A. loki* is reported both in absolute terms (black) and normalized to take into account the relative sizes of the human and archaebacterial proteomes (grey).

For mass spectrometry search algorithms, we used four current state-of-the-art programs, chosen for their distinct methods for candidate sequence selection and FDR estimation: SEQUEST HT with Percolator as part of Proteome Discoverer [16,63], Byonic [65,66], MaxQuant [67,68], and PEAKS [69]. For benchmarking purposes, we searched the experimental human K052 and negative control bacterial *E. coli* mass spectra against a concatenated database containing PGX and UniProt human databases, supplemented with negative control archaebacterial *A. loki* proteins and the common contaminant cRAP sequences. All searches were performed using identical search parameters at the FDR of less than 0.01. We assessed sensitivity based on the number of *bona fide* correct identifications, mapping to either human or common contaminant databases (Figure 4C, Supplementary table 1).

We estimated specificity from the fraction of peptides identified from *E. coli* spectra matched to human protein sequences, and experimental human spectra matched to archaebacterial *A. loki* sequences (Figure 4D). To empirically estimate FDR, we normalized the number of observed peptide-spectrum matches to account for the differences in size between human and archaebacterial proteomes.

We found that algorithms that select candidate sequences based on *de novo* sequence tags, such as Byonic and PEAKS, have a lower probability to match spectra to incorrect sequences in the database, and are thus preferred for proteogenomic analyses. We observed that with the exception of Byonic, all algorithms had comparable sensitivity. However, PEAKS also exhibited the lowest rate of incorrectly matching *E. coli* spectra to human sequences, and thus had the highest overall specificity among the compared algorithms. This finding, combined with its high sensitivity, led us to select PEAKS for integration into ProteomeGenerator as the algorithm for peptide-spectrum matching and FDR control.

### Identification of non-canonical protein isoforms using ProteomeGenerator

Compelled by the high quality of the ProteomeGenerator-constructed PGX database and genome-scale mass spectrometry, and high sensitivity and specificity of mass spectral identification by PEAKS, we used this approach to match the experimental spectra from K052 cells against either PGX or UniProt databases at the FDR of less than 0.01. Based on the same set of 2,736,597 fragmentation spectra, we observed 611,275 and 621,960 peptide-spectrum matches (22% PSM rate) when searching PGX and UniProt databases, respectively (Figure 5A, Supplementary Tables 2-3).

**Figure 5.**
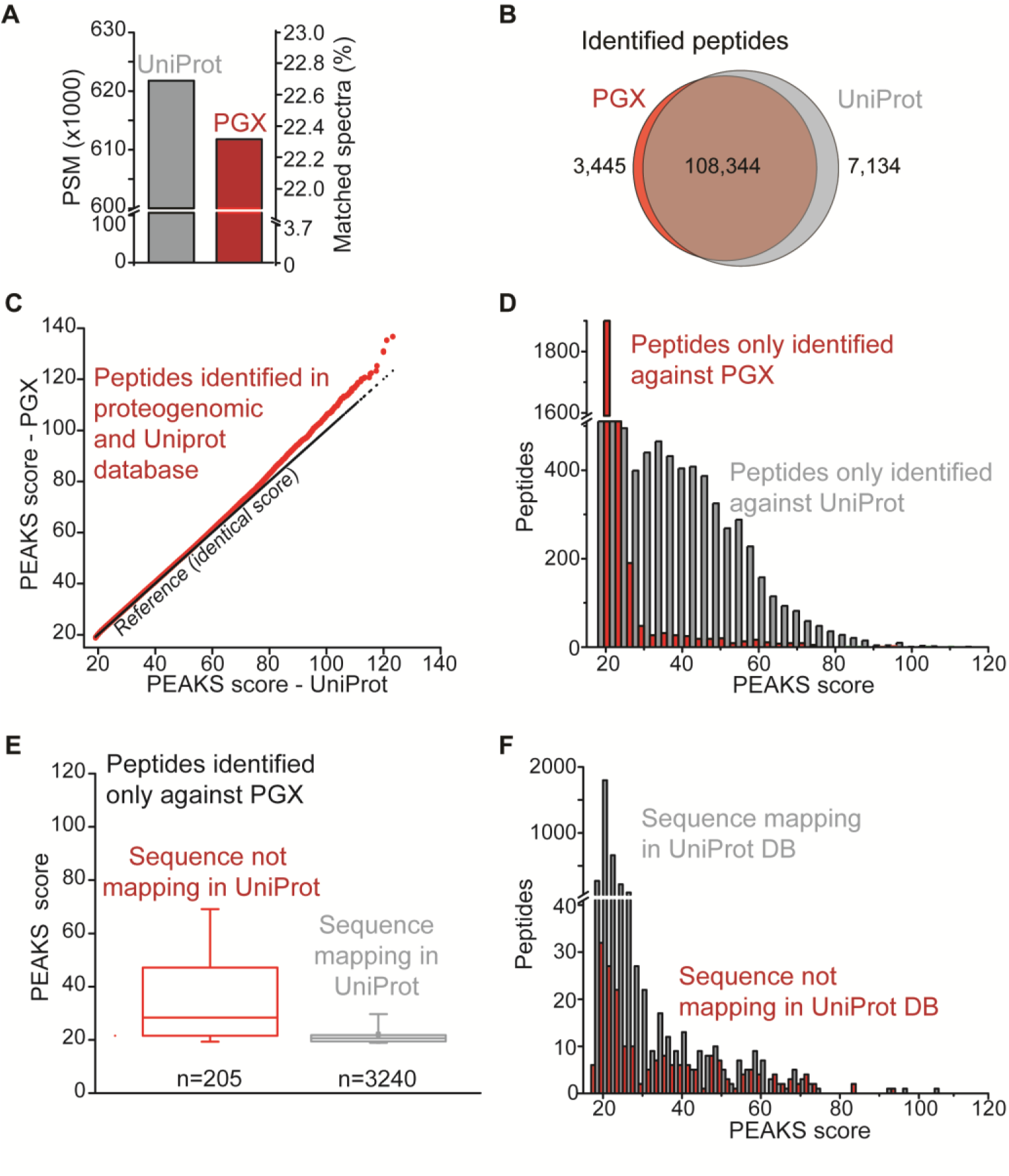
Accurate proteome discovery using statistical target-decoy matching with spectral calibration. (A) Number of peptides identified (FDR<0.01) based on matching spectra from K052 proteome against proteogenomic (PGX, red) and canonical (UniProt, grey) databases. (B) Overlap between the peptides identified in PGX (red) and UniProt (grey) databases. (C) Comparison of PEAKS scores for peptides identified in both PGX and UniProt databases. (D) PEAKS score distribution for peptides identified exclusively in PGX (red) and UniProt (grey) databases. (E) For peptides exclusively identified against the PGX database, PEAKS score distributions for peptides not mapping in UniProt (red) or present in the canonical database (grey). Boxes delimit 25th and 75th percentiles, middle line corresponds to median, whiskers correspond to 5th and 95th percentiles. (F) PEAKS score distributions for peptides identified exclusively in PGX but also mapping in UniProt (grey), or exclusively mapping in the PGX database (red).

Most of these PSMs defined peptide sequences that were shared between the two databases with 97% and 94% of the total PGX and UniProt identifications, respectively (Figure 5B), and with nearly identical confidence of identification (Figure 5C). We observed peptides uniquely identified when searching against UniProt, due to either their incomplete representation in the PGX database because of limited mRNA sequencing sensitivity, as previously described in proteogenomic analysis of HeLa cells [13], or because the greater diversity of sequences in UniProt can result in incorrect matching of homeometric peptides [76]. Importantly, the reduced size of the sample-specific PGX database led an effectively lower PSM score threshold as compared to searches against Uniprot. As a result, at identical global FDR, searches of the ProteomeGenerator-constructed PGX database were effectively more sensitive (Figure 5D–F).

We next assessed the ability of ProteomeGenerator to identify non-canonical protein isoforms. To this end, we analyzed the subset of peptides with sequences not mapping in UniProt, as prioritized based on the apparent statistical confidence of their identifications, with PEAKS peptide-spectrum matching scores greater than 50. Analysis of these sequences using BLAST indicated that the majority of them (94%) mapped to isoforms annotated in the non-reviewed section of UniProt or RefSeq (Supplementary Table 4). Notably, we identified peptides corresponding to previously un-annotated isoforms of APH (*APEH*), YB-1 (*YBX1*), and MUNC13D (*UNC13D*). These isoforms were found to be consistent with alternative splicing and intron retention for APEH and MUNC13D (Figure 6AB, Supplementary Figure 2-3).

**Figure 6.**
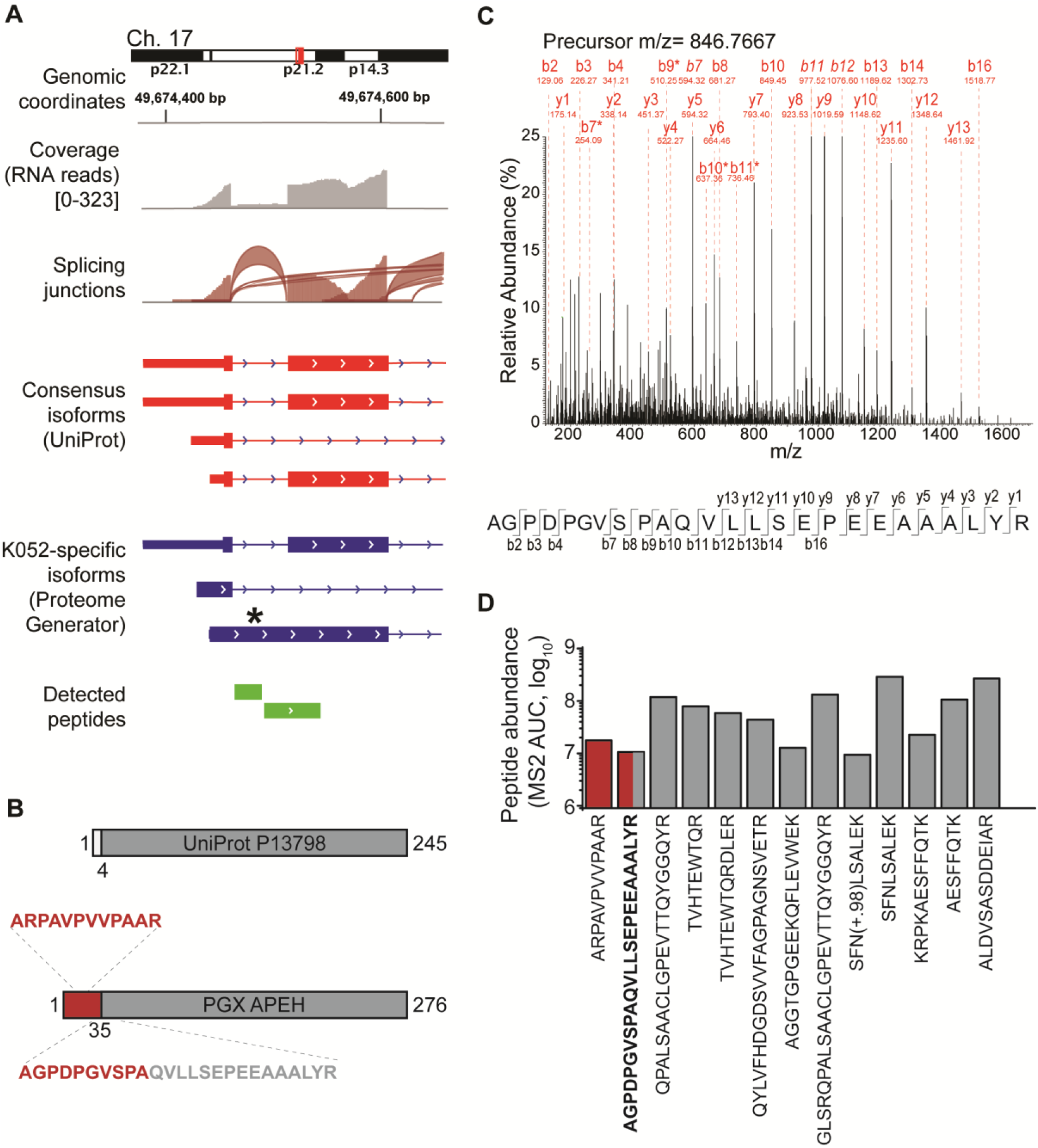
Identification of non-canonical protein isoforms using ProteomeGenerator. (A) Genome tracks of non-canonical APEH isoform generation via inclusion of an intronic sequence normally spliced in the canonical APEH isoform. (B) The K052-specific isoform of APEH contains a novel N-terminal sequence, with the splicing junction encompassed by peptide AGPDPGVSPAQVLLSEPEEAAALYR. Residues 35-276 of the protein sequences defined by ProteomeGenerator are identical to residues 4-245 of the canonical UniProt protein sequence. (C) Fragmentation spectrum of the peptide encompassing the novel splice junction, with diagnostic fragment ions and amino acid residues labeled. Italicized ion labels indicate ions with relative intensity above 25% of the maximum. A star (*) symbol denotes internal ions. (D) Peptide abundance as based on total fragment ion current for all identified APEH peptides (red: peptides from K052-specific sequence; grey: peptides from canonical sequence).

We confirmed these identifications by manual inspection of the mass fragmentation spectra (Figure 6C, Supplementary Figure 2).

In the case of APEH, the non-canonical N-terminal domain produced by intron retention was detected by mass spectrometry from two unique and independent peptides, including one spanning the non-canonical splice junction. Since no peptides corresponding to this region were detected from the canonical isoform, we used the total ion current of the fragmentation spectrum associated with this PSM to estimate the differential abundance of the novel and canonical APEH protein isoforms (Figure 6D). This analysis revealed that the identified non-canonical APEH isoform represents the majority of cellular APEH, identified specifically by ProteomeGenerator.

## DISCUSSION

Here, we introduced an analytical framework for scalable *de novo* and reference-guided assembly of sample-specific proteomic databases based on mRNA sequencing, as combined with spectral calibration high-resolution, high-accuracy mass spectrometry for discovery of non-canonical biological proteomes. We designed ProteomeGenerator to produce sample-specific proteome search databases containing only proteins that are predicted to be expressed. As a result, PGX databases have markedly reduced search space as compared to canonical reference databases such as UniProt, and consequently, exhibit more accurate control of potential false discovery by statistical database matching techniques. Since currently no methods exist for *a priori* definition of complete transcriptomes and proteomes, we confirmed the accuracy of ProteomeGenerator based on three measures: i) the set of peptides identified using ProteomeGenerator largely overlaps with that obtained by conventional matching against consensus human proteomes; ii) the novel non-canonical peptides identified by ProteomeGenerator were confirmed by high-confidence observations both at proteomic and transcriptomic levels, or were annotated in other non-reviewed or provisional shotgun databases; and iii) the accuracy of peptide-spectral matching was confirmed by stringent benchmarking of scoring functions and their calibration using spectral and sequence negative controls.

ProteomeGenerator and related approaches are limited by the accuracy of transcriptome assembly, which is dependent on mRNA sequencing quality, depth, length, and strand specificity. We anticipate that ProteomeGenerator will benefit from increasing adoption of long-read and strand-specific RNA sequencing technologies. To this end, it will also be important to further improve the sensitivity of proteome sampling by peptide mass spectrometry, such as optimizing proteome proteolysis, its chromatographic resolution, and methods for *de novo* mass spectral identification [77,78]. Lastly, ProteomeGenerator is implemented as a Snakemake workflow, enabling open, scalable, and facile discovery of sample-specific, non-canonical and neomorphic biological proteomes. This should facilitate the discovery of natural variation in cellular and tissue proteomes, which can contribute to normal tissue development and its dysregulation in human disease such as cancer.

## Author Contributions

P.C., J.T. and A.K. designed the study, JT. wrote the ProteomeGenerator computer scripts, A.Y. and O.AW. performed mRNA sequencing, P.C. and A.D. performed proteomic analysis, P.C., J.T. and A.K. performed data analyses, P.C., J.T. and A.K. wrote the manuscript with contributions from other authors.

## Acknowledgements

We thank John Phillips for constructive discussion, Nathaniel Kwok for assistance with computer scripting, Wilfred Tang for assistance with Byonic searches, and Robert Holmes for computer server support.

## Funding

This work was supported by the NIH R21 CA188881, R01 CA204396, P30 CA008748, Burroughs Wellcome Fund, Josie Robertson Investigator Program, Rita Allen Foundation, Alex’s Lemonade Stand Foundation, American Society of Hematology, and Gabrielle’s Angel Foundation (A.K.), and American Italian Cancer Foundation (P.C.). A.Y. is supported by grants from the Aplastic Anemia and MDS International Foundation and the Lauri Strauss Leukemia Foundation. A.K. is the Damon Runyon-Richard Lumsden Foundation Clinical Investigator.

## This article contains Supplemental Material

Competing Financial Interests: The authors have no competing financial interests.

---

**Supplementary Figure 1**. Subsampling analysis of mass spectrometric (A) RNA sequencing (B) dataset. Plateauing of the number of sequences detected indicates reaching of the limit of detection for the specific analytical method used, as observed in transcriptomic (B) but not in proteomic (A) datasets.

**Supplementary Figure 2**. Raw spectra for peptide AGPDPGVSPAQVLLSEPEEAA-ALYR showing (A) precursor ion (in red: monoisotopic peak) within 5 ppm from expected m/z (846.4343). (B) Original fragmentation spectrum as per Figure 6C, without intensity axis resizing.

## TABLES LEGENDS

**Supplementary Table 1**. Calibration of peptide-spectral matching using negative controls.

**Supplementary Table 2**. Peptides matched to the PGX database at FDR<0.01, using PEAKS.

**Supplementary Table 3**. Peptides matched to the UniProt database at FDR<0.01, using PEAKS.

**Supplementary Table 4**. Identified peptides with PEAKS score higher than 50 and not mapping the reference UniProt database.

This manuscript contains 38 pages, including this one.

